# ZER1 contributes to the carcinogenic activity of high-risk HPV E7 proteins

**DOI:** 10.1101/2022.07.18.500567

**Authors:** Joangela Nouel, Elizabeth A. White

## Abstract

Human Papillomavirus (HPV) E7 proteins bind to host cell proteins to facilitate virus replication. Interactions between HPV E7 and host cell proteins can also drive cancer progression. We hypothesize that HPV E7-host protein interactions specific for high-risk E7 contribute to the carcinogenic activity of high-risk HPV. The cellular protein ZER1 interacts with the E7 protein from HPV16, the genotype most frequently associated with human cancers. The HPV16 E7-ZER1 interaction is unique among HPV E7 tested to date. Other E7 proteins, even from closely related HPV genotypes, do not bind ZER1, which is a substrate specificity factor for a CUL2-RING ubiquitin ligase. In the present study, we investigated the contribution of ZER1 to the carcinogenic activity of HPV16 E7. First, we mapped the ZER1 binding site to specific residues on the C-terminus of HPV16 E7. We showed that the mutant HPV16 E7 that cannot bind ZER1 is impaired in the ability to promote the growth of primary keratinocytes. We found that ZER1 and CUL2 contribute to but are not required for HPV16 E7 to degrade RB1. Cancer dependency data shows that ZER1 is an essential gene in most HPV-positive, but not HPV-negative, cancer cell lines. Depleting ZER1 impaired the growth of primary keratinocytes expressing HPV16 E7 or HPV18 E7 and of HPV16 and HPV18-positive cervical cancer cell lines. Taken together, our work demonstrates that ZER1 contributes to HPV-mediated carcinogenesis and is essential for the growth of HPV-positive cells.

**IMPORTANCE:** HPV16 is highly carcinogenic and is the most predominant HPV genotype associated with human cancers. The mechanisms that underlie differences between high-risk HPV genotypes are currently unknown, but many of these differences are likely attributable to the activities of the oncogenic HPV proteins, including E7. The HPV E7 oncoprotein is essential for HPV-mediated carcinogenesis. A large number of HPV E7 targets have been identified. However, it is unclear which of these many interactions contribute to the carcinogenic activity of HPV E7. Here, we characterized the interaction between HPV16 E7 and the host cell protein ZER1, testing whether this genotype-specific interaction could enable the enhanced carcinogenic activity of HPV16 E7. We found that ZER1 binding contributes to the growth promoting activity of HPV16 E7 and to the growth of HPV-positive cervical cancer cells. We propose that ZER1 makes an important contribution to HPV-mediated carcinogenesis.

## INTRODUCTION

Human Papillomaviruses (HPVs) are a diverse family of small double-stranded DNA viruses with a specific tropism for keratinocytes in stratified mucosal or cutaneous epithelia. Several hundred HPV genotypes have been identified, but only up to 15 ‘high-risk’ HPVs from the genus alpha cause cancer at mucosal sites (1). The oncogenic HPVs cause almost all cases of cervical cancer and a growing number of head and neck squamous cell carcinomas (HNSCC). These and other malignancies caused by HPV infection together account for at least 4.5% of human cancer cases worldwide (2, 3).

HPV16 and HPV18 are the high-risk genotypes most commonly associated with human cancers (2). HPV16 is the most prevalent and carcinogenic genotype, causing about 60% of all cervical cancer cases and 70% of HPV-positive oropharyngeal tumors (3, 4). In contrast to high-risk HPV, ‘low-risk’ HPV genotypes cause lesions that infrequently progress to cancer. Almost all HPVs encode E6 and E7 oncoproteins that are produced throughout the papillomavirus life cycle and, for the high-risk HPVs, are the major drivers of oncogenesis. HPV E6 and E7 are small multifunctional proteins that lack enzymatic activity and alter the cellular environment largely by binding to host cellular proteins (5). Several groups have undertaken systematic approaches to identify host targets of HPV E6 and E7 (6-9).

HPV E7 proteins are about 100 amino acids long, with a disordered N-terminus and a structured C-terminus that contains two zinc binding motifs (10-12). The E7 N-terminus contains two regions that share sequence similarity with a portion of conserved region 1 (CR1) and with conserved region 2 (CR2) in the Adenovirus E1A protein (Ad E1A) (13-15). CR2 contains the LxCxE motif through which HPV E7 and Ad E1A bind to the retinoblastoma tumor suppressor (RB1) (16, 17). HPV E7 proteins bind and inactivate RB1 to release E2F transcription factors (18, 19). The disruption of the RB1/E2F complex is needed for the transcriptional activation of target genes that are essential to transition from the G1 to S phase of the cell cycle and promote cellular proliferation (20). Other conserved sequences in HPV E7 enable binding to additional host cellular proteins. These include E7 binding to the ubiquitin ligase UBR4 via sequences in CR1 and binding to the tumor suppressor PTPN14 using sequences in the C-terminus (8, 21-23). Many other host protein interactions with HPV E7 from one or more genotypes have been documented (5).

In assays of oncogenic transformation, high-risk HPV E7 have activities that are not shared by low-risk HPV E7 (24). For instance, high-risk but not low-risk HPV E7 proteins can immortalize human epithelial cells (25, 26). Although E7 from high- and low-risk HPVs can bind RB1, only high-risk E7 target RB1 for degradation (27-29). Aspects of the mechanism by which high-risk HPV E7 target RB1 for proteasome-mediated degradation have been proposed (8, 30, 31), but there are open questions regarding how high-risk HPV E7 destabilize RB1 and whether they all do so in the same way. Finally, mutational analyses of HPV E7 and studies in knockout mice show that RB1 degradation is insufficient to explain high-risk HPV E7 transforming activities (21, 32-41). We hypothesize that HPV E7-host protein interactions specific for high-risk E7 contribute to their carcinogenic activity and/or account for the increased carcinogenic potential of certain high-risk HPV genotypes.

A previous proteomic study of HPV E7-host protein interactions identified the host protein ZER1 as an interacting partner of HPV16 E7 (8). ZER1 did not bind to other closely related high-risk HPV E7 nor to HPV E7 proteins from low-risk genotypes. ZER1 belongs to the Zyg11 family of proteins and is a substrate specificity factor for a Cullin 2-RING ubiquitin ligase complex (CRL) (42). It has an N-terminal von Hippel-Lindau (VHL) box, three central leucine rich repeats (LRR), and a C-terminal armadillo-like (ARM-like) domain (42, 43). The VHL box mediates ZER1 binding to Elongin C (ELOC) and Cullin 2 (CUL2) whereas the ARM-like domain recruits substrates to the CRL (42, 43). ZER1 has been reported to specifically bind substrates that contain N-terminal glycine degrons (44). N-terminal glycine degrons are important in the quality control of N-myristoylated proteins and in eliminating protein fragments that result from apoptotic caspase cleavage (44). In addition, the N-terminal glycine degron pathway is involved in NLRP1 inflammasome activation. During enterovirus infection, the autoinhibitory NLRP1 N-terminal fragment is degraded by the ZER1/CUL2 and ZYG11B/CUL2 complexes (45). Some studies support that C-terminal sequences in HPV16 E7 recruit a CUL2-containing CRL, perhaps one that also contains ZER1, to induce the proteasomal degradation of RB1 (8, 30). However, other mutational analyses indicate that HPV16 E7 mutants that only weakly bind CUL2 retain the ability to reduce steady-state RB1 protein levels (46).

In this study, we tested the contribution of ZER1 to the carcinogenic activity of HPV16 E7. We mapped the ZER1 binding site on HPV16 E7 and used a ZER1 binding-deficient mutant of HPV16 E7 to test how ZER1 binding contributes to the growth-promoting activity of HPV16 E7. We found that primary keratinocytes expressing the HPV16 E7 ZER1 binding-deficient mutant exhibited reduced growth compared to cells expressing wild-type HPV16 E7. In a biochemical purification experiment we found that HPV16 E7, ZER1, and CUL2 co-eluted in fractions that also contained RB1. However, the majority of RB1 was still degraded when HPV16 E7 could not bind to ZER1 and CUL2, and the HPV16 E7 mutant that could not bind ZER1 could still promote the transcription of E2F target genes. Cancer gene dependency data indicated that ZER1 scored as an essential gene in nearly every HPV-positive cell line but in almost no cell lines from other cancer types. Consistent with these data, depletion of ZER1 impaired the growth of primary keratinocytes expressing either HPV16 or HPV18 E7 and reduced the anchorage-independent growth of HPV-positive cervical cancer cell lines. Our work emphasizes that ZER1 is an important contributor to the carcinogenic activity of high-risk HPV E7 and supports that ZER1 is essential for the growth of HPV-positive cancers.

## RESULTS

### HPV16 E7 potently promotes the growth of primary keratinocytes

High-risk HPV E6 and E7 are sufficient to immortalize primary keratinocytes. Keratinocyte immortalization is most efficient in the presence of high-risk E6 and E7, but high-risk E7 alone can extend keratinocyte lifespan in the short term and can promote less efficient immortalization in the longer term (24-26, 47, 48). HPV16 E6 and E7 promote cell growth more potently than other high-risk HPV oncoproteins, including HPV18 E6 and E7 (49). We tested whether, in a short-term keratinocyte growth experiment, HPV16 E7 alone had a greater growth-promoting activity than HPV18 E7. We transduced primary human foreskin keratinocytes (HFK) with retroviruses encoding high-risk HPV16 E7, HPV18 E7, or GFP as a control and tracked population doublings for about 30 days. When we monitored keratinocyte growth for several weeks, we found that HPV16 E7-expressing cells doubled more rapidly than HPV18 E7-expressing cells (Figure 1A). As expected, GFP did not promote cell growth in primary keratinocytes. HPV16 and HPV18 E7 proteins were produced at similar levels in the two HFK populations (Figure 1B). These data support that HPV16 E7 has an enhanced capacity to promote proliferation compared to HPV18 E7.

**Figure 1.**
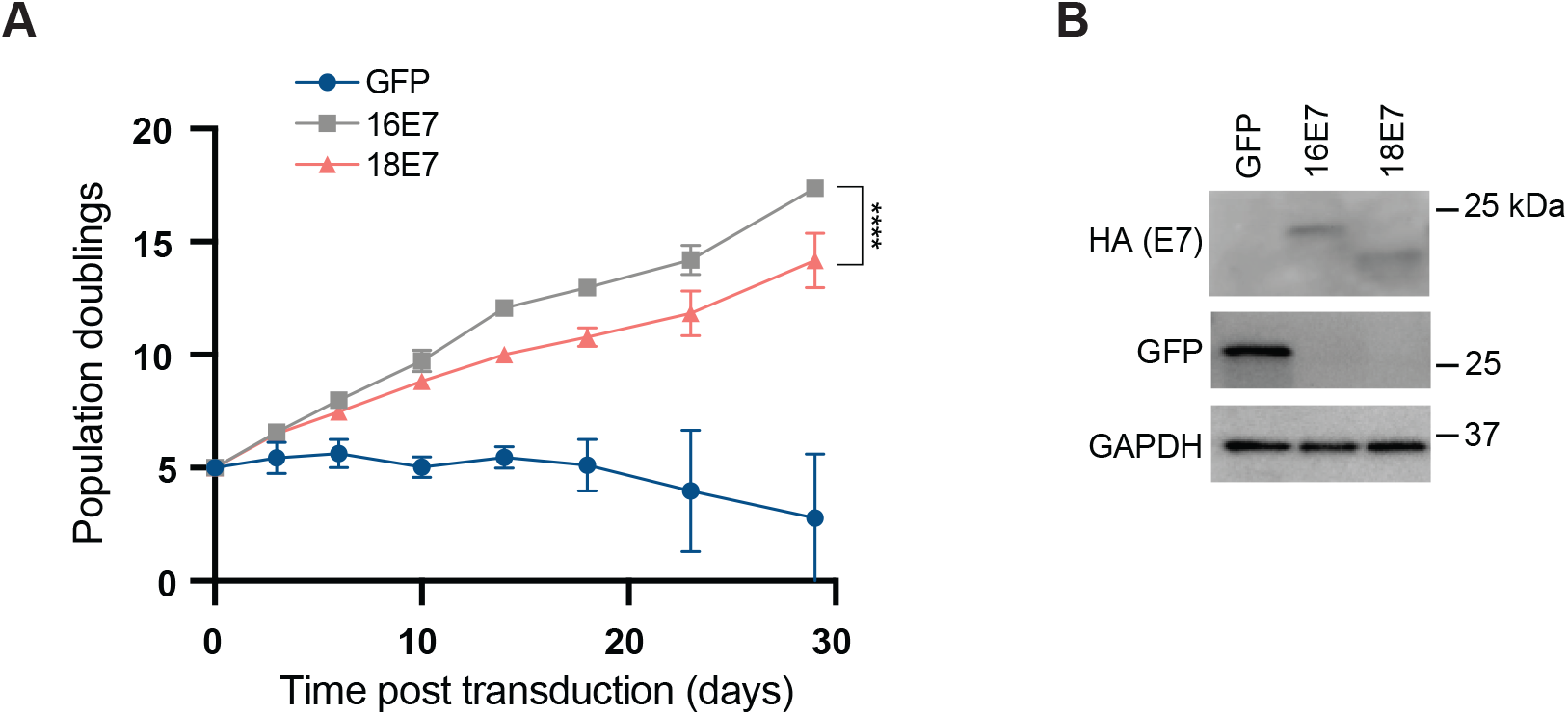
HPV16 E7 potently promotes the growth of primary keratinocytes. *(A)* Primary HFK were transduced with retroviral vectors encoding GFP or HPV E7. Population doublings were calculated based on the number of cells plated and collected at every passage. Statistical significance of the difference between HPV16 E7 and HPV18 E7 at day 29 was determined by two-way ANOVA with Sídák correction for multiple comparisons (****, p<0.0001). *(B)* Western blot of primary HFK expressing E7-HA or GFP. Cell lysates were separated by SDS/PAGE and Western blotted with antibodies to HA, GFP, and GAPDH.

### ZER1 binds to the C-terminal domain of HPV16 E7

We hypothesized that the enhanced growth-promoting activity of HPV16 E7 could be related to its unique ability to bind ZER1 (8). First, we sought to determine which region of HPV16 E7 binds ZER1. We performed a stepwise experiment to map the ZER1 binding site on HPV16 E7. Having determined that ZER1 binds to HPV16 E7 but not HPV18 E7, we created Flag-HA-tagged chimeric HPV E7 proteins in which regions of the amino and carboxyl termini were exchanged between HPV16 E7 and HPV18 E7 (Figure 2A). We transduced hTert-immortalized primary foreskin human keratinocytes (N/Tert-1) with retroviruses encoding wild-type HPV16 E7 or HPV18 E7 or the chimeric HPV E7 proteins and selected stable cell pools. Next, we performed HA immunoprecipitation (IP)-Western bIot (WB) experiments using the stable cells. As expected, ZER1 co-immunoprecipitated with HPV16 E7 but not HPV18 E7 (Figure 2B). ZER1 co-immunoprecipitated with HPV E7 chimeras B and C but not with chimera A. Chimera A contained the C-terminal domain of HPV18 E7, suggesting that the ZER1 binding site is on the C-terminus of HPV16 E7. Each chimera bound to and degraded RB1.

**Figure 2.**
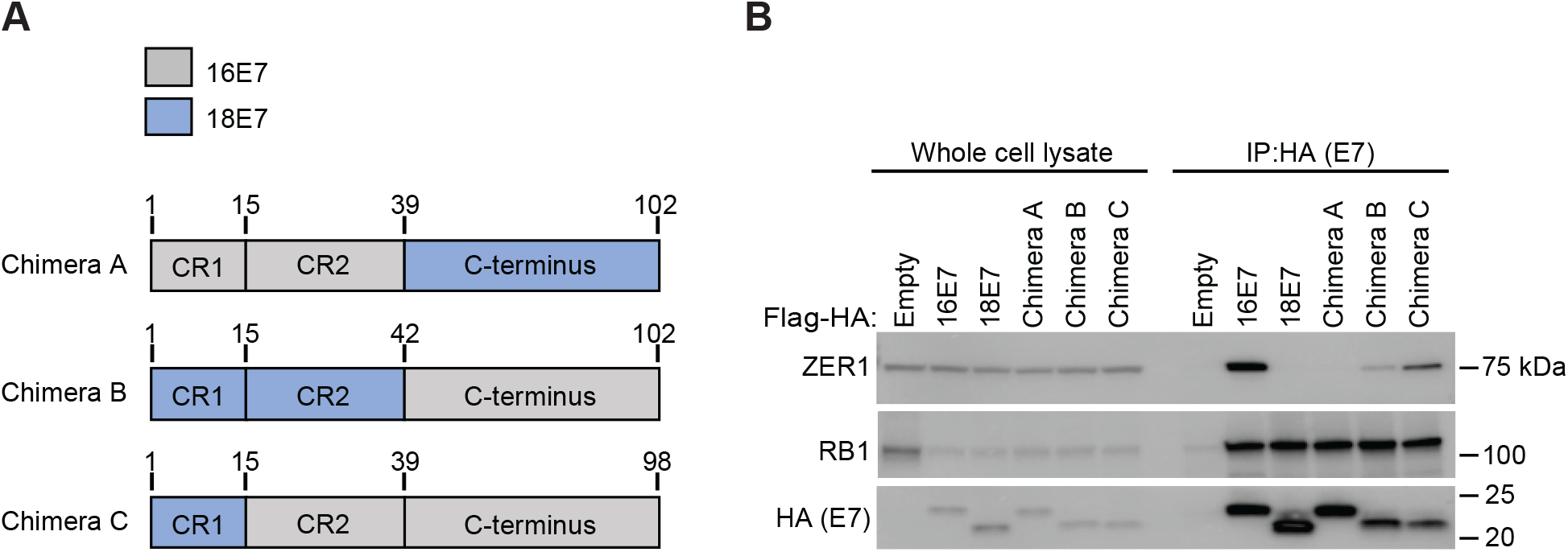
ZER1 binds to the C-terminal domain of HPV16 E7. *(A)* Schematic of HPV16-HPV18 E7 chimeras. *(B)* N/Tert-1 keratinocytes stably expressing wild-type HPV16 E7- or HPV18 E7-FlagHA, the HPV16-HPV18 E7-FlagHA chimeras, or an empty vector were subjected to anti-HA immunoprecipitation. Whole cell lysates *(left)* and immunoprecipitates *(right)* were separated by SDS/PAGE and Western blotted with antibodies to HA, ZER1, and RB1.

### ZER1 binding maps to residues E80 and D81 on the C-terminus of HPV16 E7

To determine which amino acid residues on HPV16 E7 are responsible for ZER1 binding, we performed alanine substitution mutagenesis on the C-terminus of HPV16 E7. We created a sequence alignment of several high-risk HPV E7 proteins that have been tested for their interaction with ZER1 (8), then used it to identify amino acids that are unique to HPV16 E7 compared to other E7 in the alignment (Figure 3A). The HPV16 E7-specific amino acids were mutated to alanine in two different versions of HPV16 E7 (Figure 3B). We generated N/Tert-1 cell lines that expressed the Flag-HA-tagged HPV16 E7 alanine substitution mutants and performed IP-WB analysis (Figure 3B). We found that ZER1 did not interact with HPV16 E7 mutant 1, suggesting that one or more of the amino acids that were mutated to alanine in that construct contributed to ZER1 binding (Figure 3C). Both alanine substitution mutants could bind RB1 but were modestly impaired in targeting RB1 for degradation. Next, we tested which of the five amino acids altered in mutant 1 were important for ZER1 binding, focusing on the amino acids that were predicted to be surface exposed and likely to participate in protein-protein interactions (50) (Figure 3D). We created two HA-tagged HPV16 E7 mutants, H73A (histidine changed to alanine at the 73 position) and E80A/D81A (glutamic and aspartic acid changed to alanine in positions 80 and 81), generated N/Tert-1 cells stably expressing the different forms of HPV E7, and performed IP-WB analysis. ZER1 failed to interact with HPV16 E7 E80A/D81A, suggesting that the residues responsible for ZER1 binding were E80 and D81 on the C-terminus of HPV16 E7 (Figure 3E). Since it has been reported that ZER1 can mediate the interaction between HPV16 E7 and CUL2 (8), we examined whether HPV16 E7 E80A/D81A bound CUL2. Consistent with the prediction that ZER1 bridges the interaction between CUL2 and HPV16 E7, CUL2 did not interact with HPV16 E7 E80A/D81A (Figure 3E). Each HPV16 E7 mutant retained the ability to bind and degrade RB1. We conclude that ZER1 binds amino acids E80 and D81 on the C-terminus of HPV16 E7. Mutating HPV16 E7 E80 and D81 also prevented CUL2 binding.

**Figure 3.**
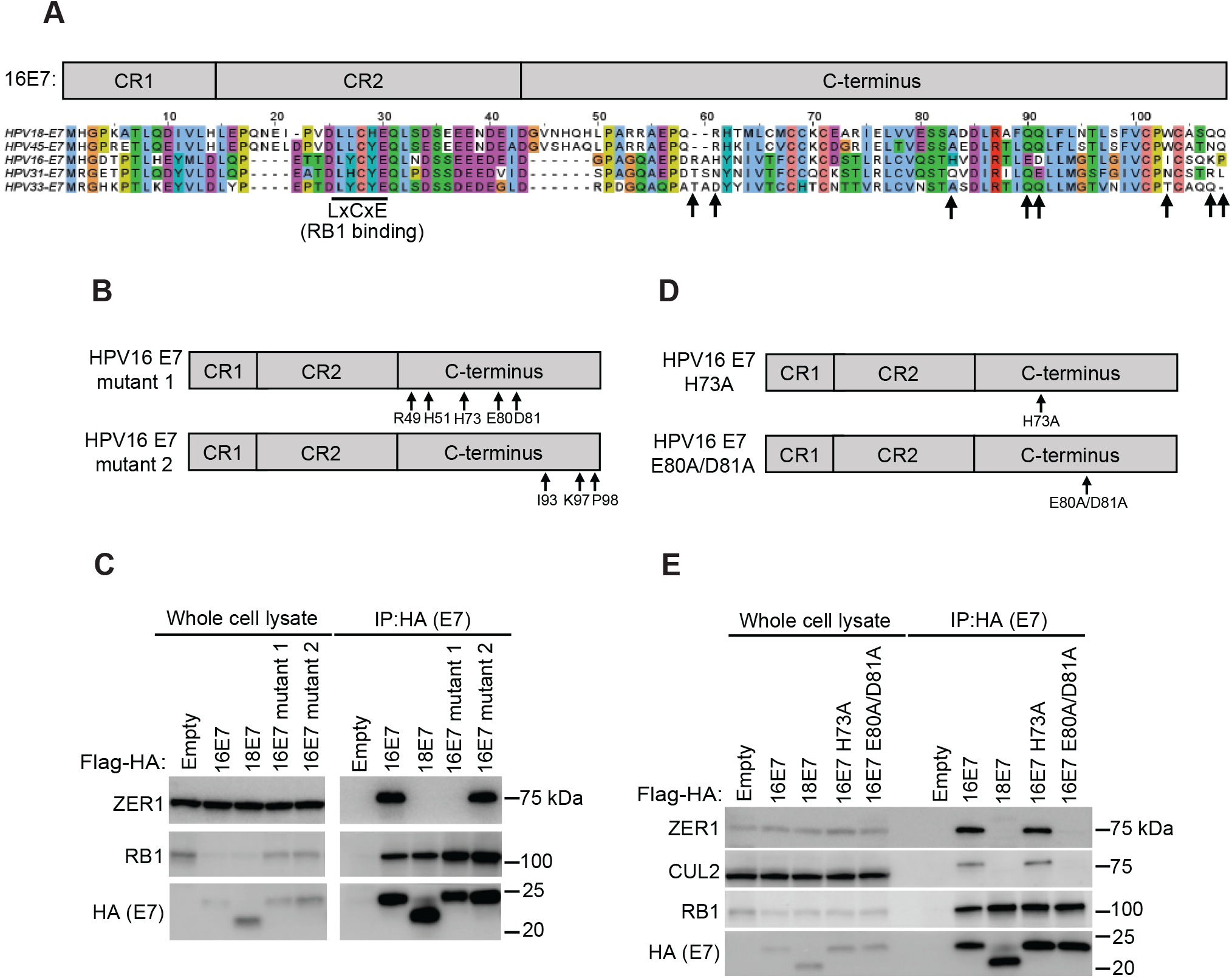
ZER1 binding maps to residues E80 and D81 on the C-terminal domain of HPV16 E7. *(A)* Protein sequence alignment of high-risk HPV E7 proteins tested for ZER1 interaction in (8). Sequences were aligned using ClustalWS and amino acids colored according to the ClustalX color scheme. Arrows indicate amino acid residues unique to HPV16 E7. The LxCxE motif responsible for RB1 binding is underlined with a black bar. *(B)* Schematic of HPV16 E7 mutants. Amino acids indicated by the arrows were substituted with alanine residues. *(C)* N/Tert-1 cells stably expressing Flag-HA-tagged wild-type HPV16 E7, HPV18 E7, the HPV16 E7 mutants, or an empty vector were subjected to anti-HA immunoprecipitation. Whole cell lysates *(left)* and immunoprecipitates *(right)* were separated by SDS/PAGE and Western blotted with antibodies to HA, ZER1, and RB1. *(D)* Schematic of additional HPV16 E7 mutants. Amino acids indicated by the arrows were substituted with alanine residues. *(E)* N/Tert-1 cells stably expressing Flag-HA-tagged wild-type HPV16 E7, HPV18 E7, the HPV16 E7 mutants, or an empty vector were subjecteded to anti-HA immunoprecipitation. Whole cell lysates and immunoprecipitates were separated by SDS/PAGE and Western blotted with antibodies to HA, ZER1, CUL2, and RB1.

### HPV16 E7 E80A/D81A is impaired in the ability to promote growth of primary keratinocytes

To test whether ZER1 binding contributed to the ability of HPV16 E7 to promote cell growth, we initiated growth curve experiments in primary HFK. PTPN14 is a tumor suppressor that we found restricts cell growth by promoting differentiation (51). Conserved amino acids in the HPV E7 C-terminus enable PTPN14 binding and degradation (52). In the N/Tert-1 cell lines expressing HPV16 E7, HPV16 E7 E80A/D81A, or an empty vector control, we observed that steady-state PTPN14 protein levels were reduced in wild-type HPV16 E7 cells but not in HPV16 E7 E80A/D81A-expressing cells (Figure 4A). To control for any effects of PTPN14 degradation on the growth of cells expressing wild-type HPV16 E7 versus HPV16 E7 E80A/D81A, we conducted keratinocyte growth experiments in the presence and absence of CRISPR/Cas9 constructs targeting PTPN14. We transduced primary HFK with retroviruses encoding wild-type HPV16 E7 or the ZER1 binding-deficient mutant HPV16 E7 E80A/D81A plus LentiCRISPRv2 lentiviruses encoding PTPN14-targeting sgRNAs or non-targeting controls. Wild-type HPV16 E7 efficiently promoted the growth of primary HFK, with only a modest additional growth increase in cells depleted of PTPN14 (Figure 4B). In contrast, HPV16 E7 E80A/D81A was impaired in its ability to promote keratinocyte growth. Similar to the effects of PTPN14 knockout in the wild-type HPV16 E7 cells, PTPN14 knockout slightly restored growth of primary keratinocytes expressing HPV16 E7 E80A/D81A. These data support that ZER1 binding contributes to the growth-promoting activity of HPV16 E7 in a manner that is largely independent of PTPN14 binding and degradation.

**Figure 4.**
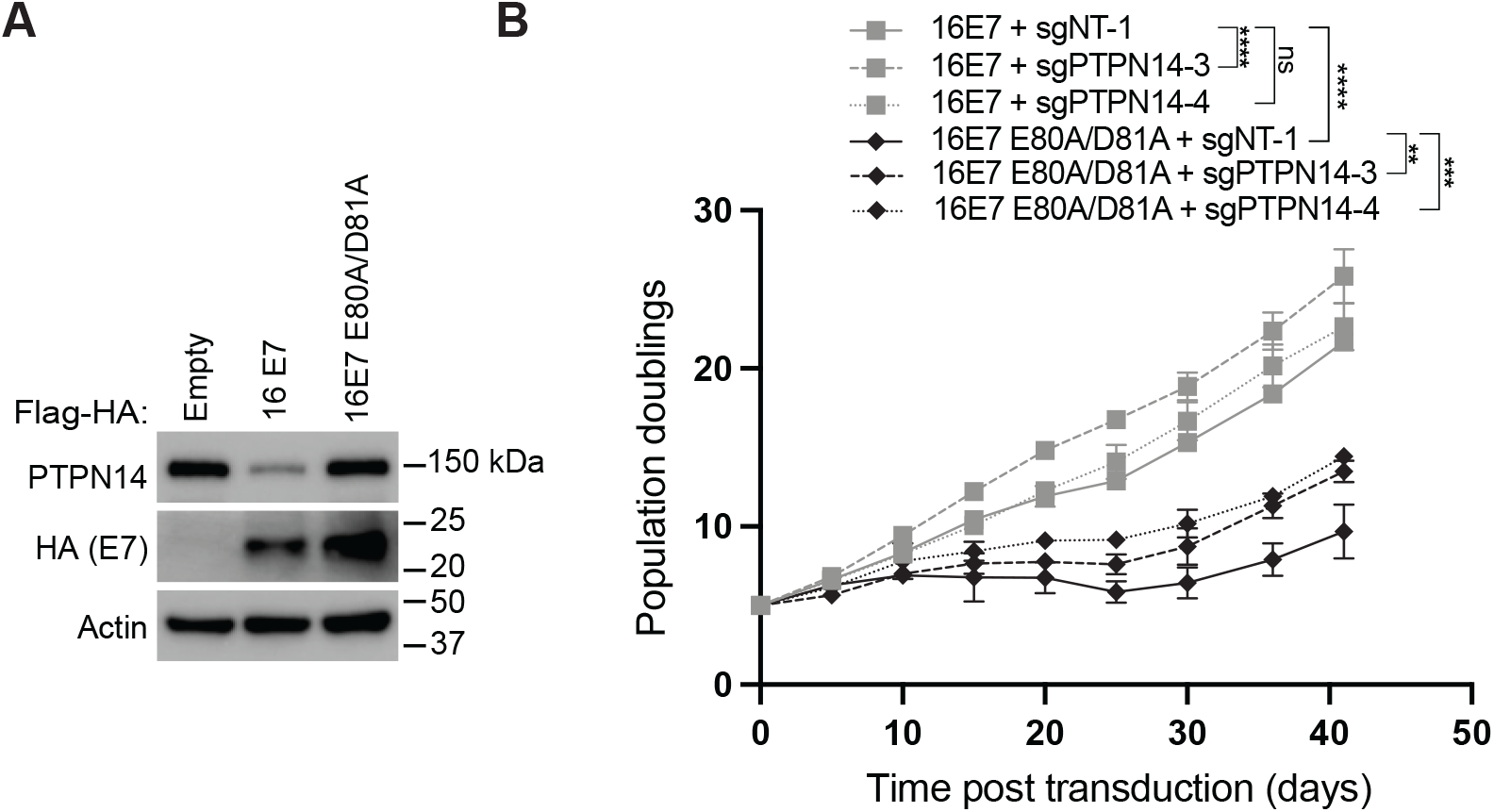
HPV16 E7 E80A/D81A impairs the growth of primary keratinocytes. *(A)* Western blot of N/Tert-1 keratinocytes expressing Flag-HA-tagged HPV16 E7, HPV16 E7 E80A/D81A or an empty vector. Cell lysates were separated by SDS/PAGE and Western blotted with antibodies to HA, PTPN14, and actin. *(B)* Primary HFK were transduced with retroviral vectors encoding HPV E7 and LentiCRISPRv2 vectors encoding PTPN14-targeting or nontargeting sgRNAs. Population doublings were calculated based on the number of cells plated and collected at every passage. Statistical significance of the difference between HPV16 E7 and HPV16 E7 E80A/D81A non-targeting control and PTPN14-targeting sgRNA conditions at day 41 was determined by two-way ANOVA with Sídák correction for multiple comparisons (ns, not significant; **, p<0.005; ***, p<0.0005; ****, p<0.0001).

### HPV16 E7, ZER1, and CUL2 contribute to RB1 destabilization

Next, we used gel filtration chromatography to further characterize the protein complex that contains HPV16 E7 and ZER1. Lysates of N/Tert-1 cells expressing wild-type HPV16 E7 were separated into 500ul fractions on a Superose 6 column. Flag-HA-tagged HPV16 E7 was immunoprecipitated from the even numbered fractions and immunoprecipitated lysates were analyzed by Western blotting. HPV16 E7 eluted from the column over a range of fractions containing protein complexes of estimated sizes from 200-2000 kDa (Figure 5A). RB1 eluted over a similar range of fractions. Peak elution of HPV16 E7 and of RB1 occurred in two sets of fractions, one around fraction 10-12 and the other around fraction 18. The fractions representing higher molecular weight complexes contained E7, UBR4, and RB1. Fractions 18-20 contained E7, ZER1, CUL2, and RB1. Previously we reported that CUL2 co-immunoprecipitates with HPV16 E7 only in the presence of ZER1 (8). Consistent with earlier reports (30), these data support that 16E7-ZER1-CUL2 are present in a multiprotein complex (Figure 5B). Some, but not all, of the RB1 that co-elutes with HPV16 E7 is in the fractions containing CUL2 and ZER1.

**Figure 5.**
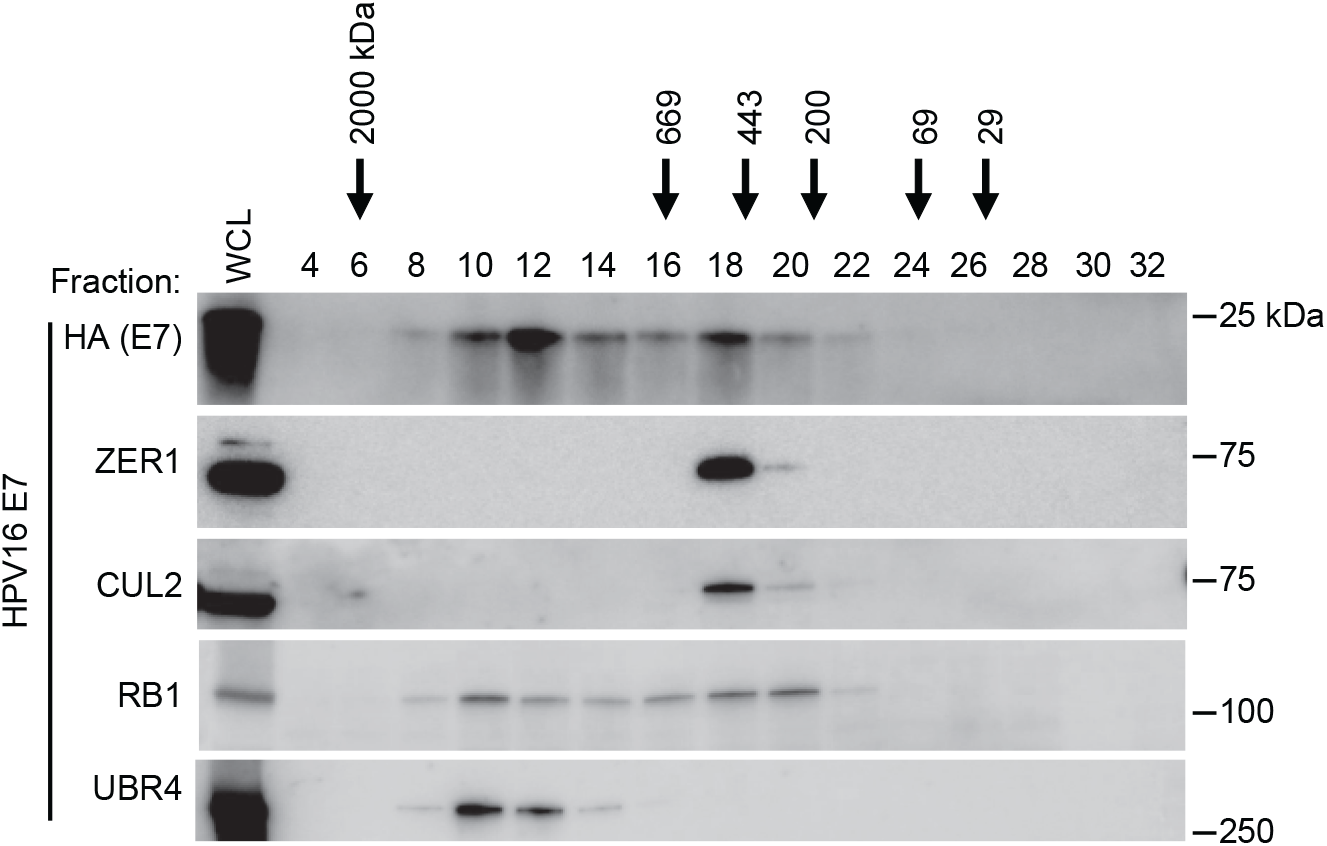
HPV16 E7 co-fractionates with CUL2 and ZER1. N/Tert-1 keratinocytes stably expressing wild-type HPV16 E7-Flag-HA were fractionated on a Superose 6 column. Five hundred-microliter fractions were collected and the even numbered fractions were subjected to anti-HA immunoprecipitation. The unfractionated whole cell lysate (WCL) and immunoprecipitates were separated by SDS/PAGE and Western blotted with antibodies to HA, ZER1, CUL2, RB1, and UBR4.

HPV16 E7 targets hypo-phosphorylated RB1 for proteasome-mediated degradation (29, 53). Having already assessed the steady-state levels of RB1 in N/Tert-1 keratinocytes expressing wild-type or mutant HPV16 E7 (Figure 3), we next used an antibody that recognizes the hypo-phosphorylated form of RB1. In this experiment we included N/Tert-1 keratinocytes that stably express HPV16 E7 ΔDLYC (does not bind RB1). As expected, RB1 levels were reduced in cells expressing HPV16 E7 but not in cells expressing HPV16 E7 ΔDLYC (Figure 6A). Compared to wild-type HPV16 E7 or to HPV16 E7 ΔDLYC cells, HPV16 E7 E80A/D81A mutant cells contained intermediate levels of hypo-phosphorylated RB1.

**Figure 6.**
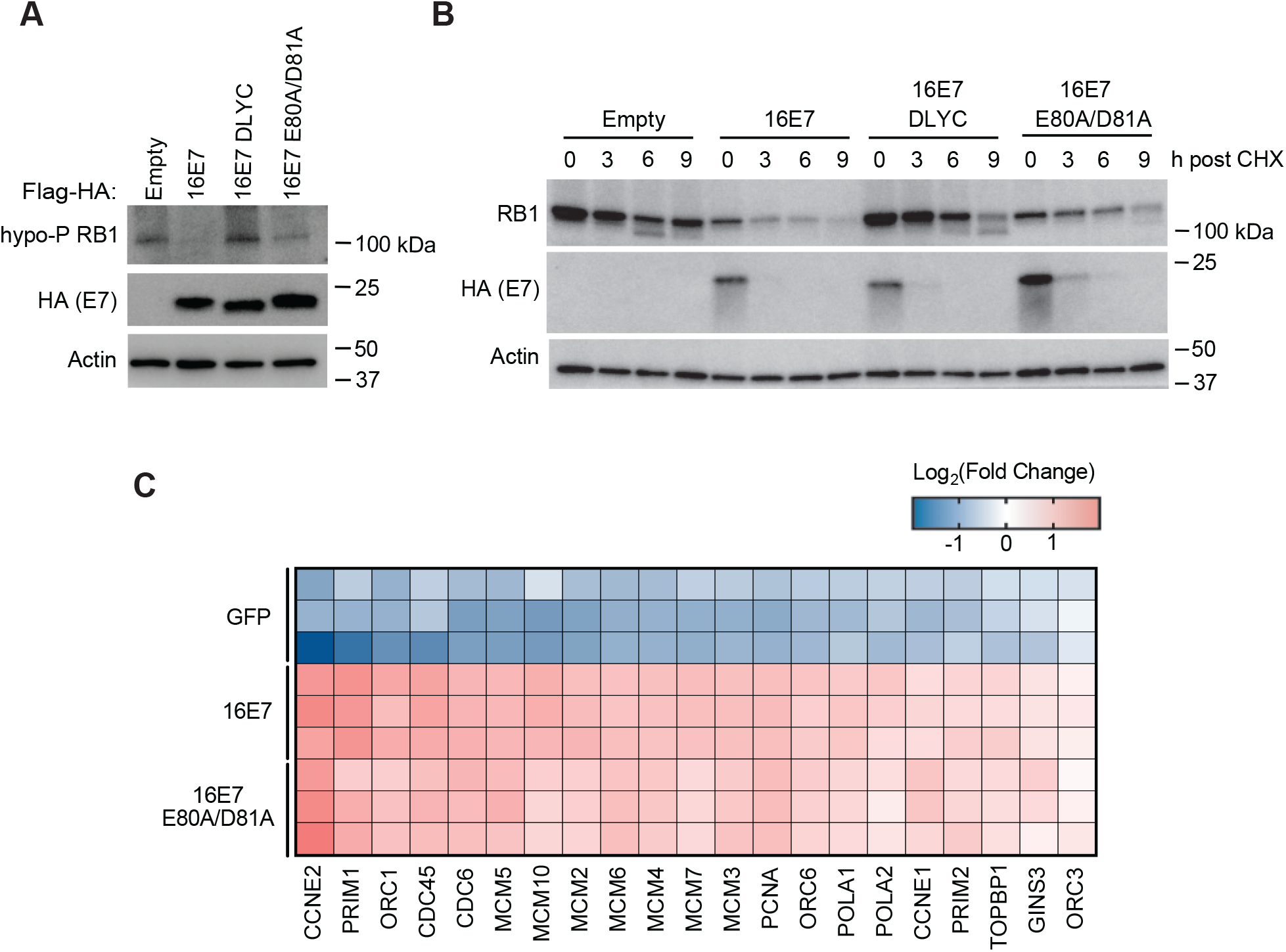
HPV16 E7 E80A/D81A prolongs the half-life of RB1. *(A)* Western blot of N/Tert-1 keratinocytes stably expressing empty vector, Flag-HA-tagged wild-type HPV16 E7, or the HPV16 E7 mutants DLYC and E80A/D81A. Protein lysates were separated by SDS/PAGE and Western blotted with antibodies to the hypo-phosphorylated form of RB1, HA, and actin. *(B)* N/Tert-1 cells stably expressing empty vector, Flag-HA-tagged wild-type HPV16 E7, or the HPV16 E7 mutants DLYC and E80A/D81A were treated with 40 ug/mL cycloheximide (CHX) and harvested at the indicated time points. Protein lysates were separated by SDS/PAGE and Western blotted with antibodies to RB1, HA, and actin. The experiment was repeated three times and a representative image is shown. *(C)* Primary HFK were transduced with retroviral vectors encoding HPV16 E7, HPV16 E7 E80A/D81A, or GFP as a control. Poly(A)-selected RNA was analyzed by RNA-seq. Heat map showing the expression patterns of selected DNA replication initiation genes.

To determine how binding to ZER1 affected the ability of HPV16 E7 to destabilize RB1, cells were treated with cycloheximide to halt *de novo* protein synthesis and harvested at the indicated time points. The half-life of RB1 decreased from 25.5 hours in vector control cells to 5.3 hours in cells expressing wild-type HPV16 E7 (Figure 6B, Supplemental figure 1). In the presence of HPV16 E7 E80A/D81A the half-life of RB1 was 8.6 hours, representing a modest increase over its half-life in wild-type HPV16 E7 cells. In contrast, RB1 was relatively stable in HPV16 E7 ΔDLYC cells, with a half-life of 15.6 hours. Similar to hypo-phosphorylated RB1 levels in these cell lines, total RB1 levels at the beginning of the experiment were highest in vector control and HPV16 E7 ΔDLYC cells, lowest in HPV16 E7 cells, and modestly restored in HPV16 E7 E80A/D81A cells. These data support that CUL2 and ZER1 contribute to, but are not the only factors responsible for, RB1 degradation by HPV16 E7.

Since the protein levels of RB1 were slightly increased in HPV16 E7 E80A/D81A-expressing cells, we wanted to determine whether the increase in RB1 levels was associated with reduced expression of E2F target genes. Reduced E2F target gene expression could account for the impaired growth phenotype in HPV16 E7 E80A/D81A-expressing cells. We sequenced polyA-selected RNA from HFK transduced with GFP, wild-type HPV16 E7, or HPV16 E7 E80A/D81A. We selected Gene Ontology (GO) term 0006270: DNA replication initiation as a representative list of E2F target genes (54). Compared to GFP control cells, E2F target genes were upregulated in cells expressing either wild-type HPV16 E7 or HPV16 E7 E80A/D81A (Figure 6C). Although RB1 protein levels were slightly increased in HPV16 E7 E80A/D81A-expressing cells, the HPV16 E7 mutant retained its ability to induce the expression of E2F target genes, thereby promoting cell cycle entry and DNA replication.

### ZER1 is essential in HPV-positive cancer cell lines

The Cancer Dependency Map (DepMap) project uses genome-wide loss-of-function screens to determine which genes are essential in hundreds of different cancer cell lines (55, 56). Gene essentiality is quantified using the Chronos dependency score, which is used to assess fitness effects in CRISPR knockout screens. Nonessential genes have a Chronos score of 0, whereas a Chronos score of −1 corresponds to the median of all common essential genes (57) and indicates that a gene is likely to be essential in a given cell line. We downloaded the ZER1 Chronos dependency scores for several hundred cancer cell lines from the DepMap website (https://depmap.org/). Nearly all of the cell lines in which ZER1 is predicted to be an essential gene were from cancers caused by HPV (Supplemental Figure 2). We graphed the data from cervical, head and neck, and endometrial/uterine cancer cell lines, finding that the ZER1 Chronos scores ranged from −0.5 to −1 for most of the HPV-positive but not the HPV-negative cell lines (Figure 7). The HPV-positive cell lines contained DNA from one of several high-risk HPV types including HPV16, 18, 31 and 56. These data from the DepMap project CRISPR screens indicate that ZER1 is essential for the viability of HPV-positive cancer cell lines.

**Figure 7.**
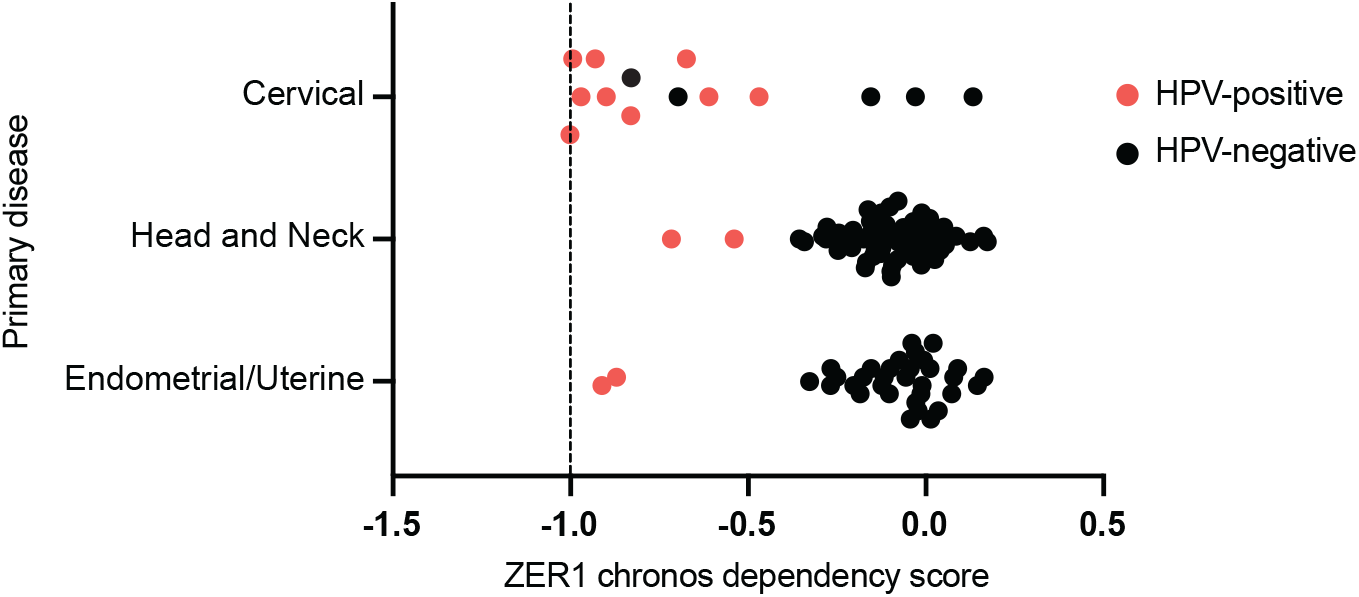
HPV-positive cells depend on ZER1 to survive. ZER1 chronos dependency scores from cervical, head and neck, and uterine cancer cell lines were downloaded from the DepMap website (https://depmap.org/). Each dot represents a cell line. A score of 0 denotes that a gene is nonessential whereas a score of −1 corresponds to the median of all common essential genes.

### ZER1 knockout impairs the growth of high-risk HPV E7-expressing keratinocytes

The gene essentiality data led us to hypothesize that ZER1 contributes to the carcinogenic activity of high-risk HPV besides HPV16. We next sought to determine whether ZER1 contributed to the growth promoting activity of either HPV16 E7 or HPV18 E7. To do so, we first established tools for CRISPR knockout of ZER1. In our initial IP-WB experiments, we determined that the ZER1 antibody specifically detected ZER1 protein after co-immunoprecipitation with HA-E7. However, in a whole cell lysate, the endogenous ZER1 band was obscured by a co-migrating background band. Therefore, we used N/Tert-1 keratinocytes stably expressing HA-ZER1 to validate LentiCRISPRv2 constructs targeting ZER1 (Figure 8A). Then, we transduced primary HFK with retroviruses encoding HPV16 E7 or HPV18 E7 and with LentiCRISPRv2 vectors encoding ZER1-targeting or non-targeting sgRNAs. We grew the cells and tracked population doublings for about 40 days. As expected, HPV16 E7 efficiently promoted the growth of primary HFK, but ZER1 knockout profoundly impaired the growth of HFK-HPV16 E7 (Figure 8B). Although ZER1 knockout impaired the growth of both HPV16 E7 and HPV18 E7 expressing cells, the growth defect was most pronounced in the HPV16 E7 cells. We conclude that ZER1 inactivation impairs the growth of HPV E7-expressing keratinocytes.

**Figure 8.**
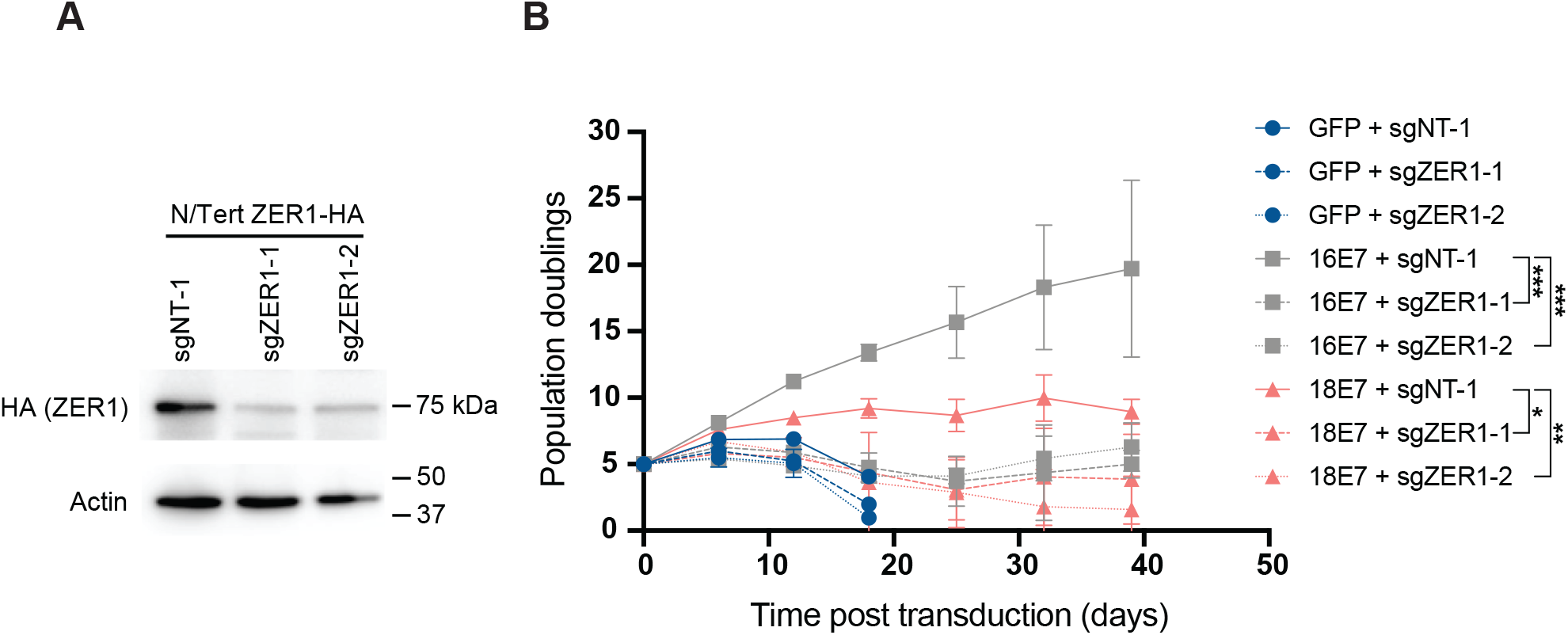
ZER1 knockout impairs the growth of HPV E7-expressing keratinocytes. *(A)* N/Tert-1 cells that stably express HA-tagged ZER1 were transduced with LentiCRISPRv2 vectors encoding ZER1-targeting or nontargeting sgRNAs. *(B)* Primary HFK were transduced with retroviral vectors encoding GFP or HPV E7 and lentiviral vectors encoding SpCas9 and ZER1-targeting or nontargeting sgRNAs. Population doublings were calculated based on the number of cells plated and collected at every passage. Statistical significance of the difference between HPV16 and HPV18 E7 ZER1 knockout conditions compared to the non-targeting control at day 39 was determined by two-way ANOVA with Sídák correction for multiple comparisons (*, p<0.05; **, p<0.01; ***, p<0.0005).

### ZER1 contributes to the anchorage-independent growth of HPV-positive cervical cancer cell lines

Next, we wanted to determine whether ZER1 was required for the anchorage-independent growth of HPV-positive cancer cell lines. Given our limited ability to detect endogenous ZER1 in whole-cell lysates, we confirmed that ZER1 is expressed in HPV-positive and HPV-negative cervical cancer cell lines by using HPV16 E7 as a bait in co-immunoprecipitation experiments. We transduced SiHa and Caski (HPV16-positive), HeLa (HPV18-positive) and C33A (HPV-negative) cells with a retrovirus encoding Flag-HA-tagged HPV16 E7 and performed IP-WB analysis. ZER1 interacted with HPV16 E7 in all cervical cancer cell lines, demonstrating that ZER1 is expressed and can bind to HPV16 E7 in each cell line (Figure 9A). Then, we performed soft agar assays to measure the anchorage independent growth of HPV-positive cervical cancer cells. We transduced SiHa, HeLa, and C33A cells with LentiCRISPRv2 constructs encoding ZER1-targeting or non-targeting sgRNAs. Soft agar assays using the modified cell lines were incubated for 3 weeks. At the assay endpoint an average of 2,000 colonies were counted in the non-targeting control conditions in SiHa and HeLa cells. ZER1 knockout caused a 10-150-fold decrease in colony formation in SiHa cells and a 6-fold decrease in Hela cells (Figures 9B and 9C). In C33A cells, ZER1 knockout only decreased colony formation by about 50% (Figure 9D). These data show that ZER1 is required for the anchorage independent growth of HPV-positive cervical cancer cells.

**Figure 9.**
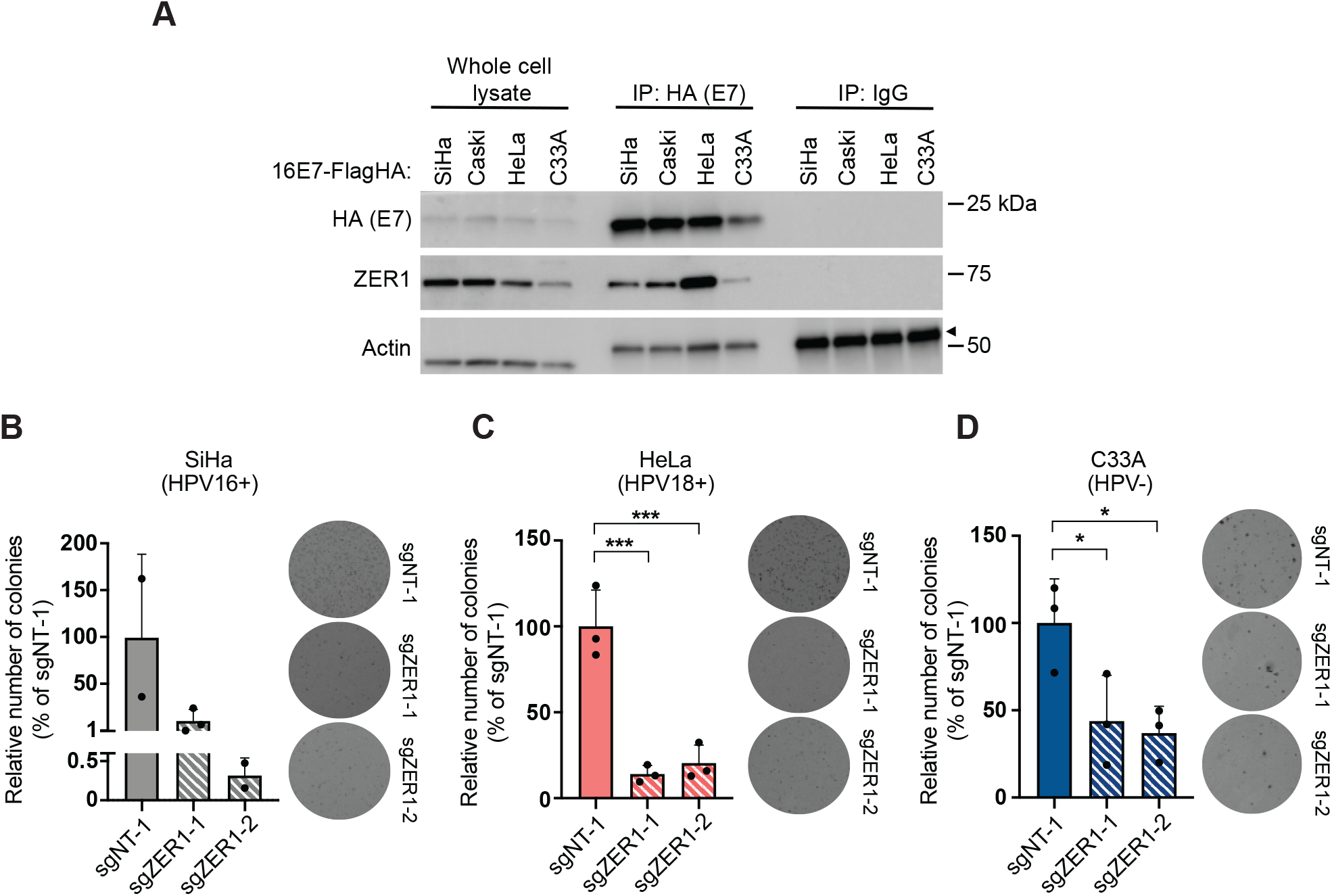
ZER1 contributes to the anchorage-independent growth of HPV-positive cervical cancer cell lines. *(A)* SiHa, Caski, HeLa, and C33A cells stably expressing wild-type HPV16 E7-Flag-HA were subjecteded to immunoprecipitation with anti-HA or control IgG. Whole cell lysates *(left)* and immunoprecipitates *(right)* were separated by SDS/PAGE and Western blotted with antibodies to HA, ZER1, and actin. Arrow indicates the heavy chain. *(B)* SiHa, *(C)* HeLa, and *(D)* C33A cell lines were transduced with lentiviral vectors encoding SpCas9 and ZER1-targeting or nontargeting sgRNAs. Cells were plated in triplicate for each condition and incubated at 37°C for 3 weeks. Plates were photographed and colonies were quantified using ImageJ. Graphs show the number of colonies relative to the non-targeting control condition for each cell line. Graphs indicate mean +/- standard deviation and display individual data points from each experiment. Statistical significance of the non-targeting control condition compared to the ZER1 knockout conditions was determined by ANOVA with the Holm-Sídák correction for multiple comparisons (*, p<0.05; ***, p<0.001).

## DISCUSSION

Although many HPV E7-host protein-protein interactions have been identified, the biological role of most of them remains to be elucidated. In addition, it is unclear which of these interactions contribute to high-risk HPV E7 carcinogenicity. One well-characterized activity of high-risk HPV E7 proteins is to bind and degrade RB1 to promote cell cycle re-entry in differentiated keratinocytes. Several HPV E7 also recruit the ubiquitin ligase UBR4 to target PTPN14 for degradation, thereby repressing keratinocyte differentiation and promoting carcinogenesis (22, 58). In contrast to RB1, PTPN14, and UBR4, which interact with HPV E7 proteins from diverse genotypes, other host proteins bind to specific HPV E7 proteins. For example, of the HPV E7 tested so far, only HPV16 E7 can interact with the host protein ZER1 (8). Compared to other high-risk HPV oncoproteins, HPV16 E6 and E7 most potently induce immortalization and promote the growth of primary keratinocytes (49). We sought to determine whether ZER1 contributes to the enhanced carcinogenic activity of HPV16 E7.

First, we used a systematic mutagenesis strategy to determine that ZER1 binds to amino acid residues E80 and D81 on the C-terminus of HPV16 E7 (Figures 2 and 3). Compared to wild-type HPV16 E7, the HPV16 E7 mutant that cannot bind ZER1 is impaired in promoting keratinocyte growth (Figure 4). These data are consistent with the idea that ZER1 binding contributes to the growth-promoting activity of HPV16 E7. HPV16 E7 has been proposed to engage a CUL2^ZER1^ ligase complex to target RB1 for degradation, and so we tested whether the inability to degrade RB1 could account for the growth defect exhibited by the HPV16 E7 ZER1 binding-deficient mutant. In gel filtration experiments, we found that HPV16 E7, ZER1, and CUL2 co-eluted together with some RB1, but that HPV16 E7 and RB1 also exist in fractions separate from CUL2/ZER1 (Figure 5). The levels of hypo-phopshorylated RB1 and the half-life of RB1 were only modestly increased in HPV16 E7 mutant cells compared to wild-type controls (Figures 6A and B) and the HPV16 E7 ZER1 binding-deficient mutant retained the ability to activate E2F target gene expression (Figure 6C). Taken together, these data support that only some RB1 is targeted for degradation by HPV16 E7/CUL2^ZER1^. Cancer dependency data suggests that ZER1 is essential in many HPV-positive cervical cancer cell lines, even those transformed by HPV genotypes other than HPV16 (Figure 7). Our data support that ZER1 is important for the growth-promoting activity of several HPV E7, showing that ZER1 depletion reduced the growth of both HPV16 E7 and HPV18 E7-expressing keratinocytes (Figure 8). Depleting ZER1 also reduced the anchorage independent growth of HPV-positive cervical cancer cells (Figure 9).

Because RB1 destabilization is thought to be an important carcinogenic activity of high-risk HPV E7, many years of research have sought to determine how oncogenic HPV E7 inactivate RB1. In proliferating cells, RB1 has a long half-life (27). Previous work suggested that HPV16 E7 recruits the CUL2^ZER1^ E3 ligase to target RB1 for degradation (8, 30). Our new data indicate that HPV16 E7/CUL2^ZER1^ contributes to a reduction in hypo-phosphorylated RB1 protein levels, but that most HPV16 E7/RB1-containing complexes do not contain CUL2^ZER1^. Furthermore, even the HPV16 E7 ZER1 binding-deficient mutant can significantly reduce RB1 levels. These data remain consistent with earlier reports, which showed that knocking down ZER1 or CUL2 with siRNAs restored some, but likely not all, hypo-phosphorylated RB1 protein in HPV16 E7-expressing cells (8, 30). Our finding that only some HPV16 E7/RB1 is complexed with CUL2^ZER1^ may result from the fact that we fractionated HPV16 E7-containing complexes over a broader range of molecular weights than in a previous report (30). Another study observed that HPV16 E7 mutants with altered C-terminal, surface exposed amino acids were still able to bind CUL2 and degrade RB1 and concluded that CUL2 could not facilitate RB1 degradation (46). That study did not examine ZER1, but did include HPV16 E7 E80K/D81K, a mutant similar to the one we described here. One difference between that report and ours is that our analyses tested endogenous RB1 levels in human keratinocytes, whereas the earlier study used overexpressed, tagged RB1. It is possible that a higher initial level of RB1 in the overexpression study limited the authors’ ability to observe an effect of HPV E7/CUL2 on RB1 levels. Overall, we propose that CUL2^ZER1^ is recruited by HPV16 E7 to target some RB1 for degradation, but that other activities of high-risk HPV E7 also destabilize RB1. We hypothesize that there is a conserved mechanism used by high-risk HPV E7 proteins to target RB1 for degradation and that the molecular basis of such a mechanism remains to be identified.

Although the HPV16 E7 ZER1 binding mutant upregulated E2F target genes, its ability to promote keratinocyte growth was significantly impaired. We therefore continued to investigate the importance of ZER1 in keratinocyte growth and in cancer cell growth. One activity of viral oncoproteins, including HPV E7, is to target host proteins for degradation. However, it is unlikely that HPV16 E7 is reducing ZER1 levels, post-translationally or otherwise. In the RNA-seq experiment, *ZER1* was expressed in the presence and absence of HPV16 E7. In our immunoprecipitation experiments, ZER1 protein was readily detectable in the cervical cancer cell lines (Figure 9A). The observations that ZER1 knockout limited cell growth in the keratinocyte proliferation assay and in assays of anchorage independent growth also support that E7-expressing cells contain functional ZER1. ZER1 binds specifically to HPV16 E7. We were therefore surprised that in our experiments and in a publicly available dataset, the requirement for ZER1 was not specific to HPV16-positive cells. Cancer dependency data emphasized that ZER1 is an essential gene in HPV-positive cancer cells transformed with several high-risk HPV genotypes, but not in most other cancer cell lines (Supplemental figure 2), highlighting that there are essential genes specific to cancers caused by HPV. It remains an open question how both binding to ZER1 by HPV16 E7 and the presence of ZER1 could be required for HPV-mediated carcinogenesis.

Our results emphasize the importance of future studies on HPV and ZER1. Most existing data on ZER1 is related to its function as a substrate specificity factor of a CUL2 E3 ligase complex. We therefore favor the model that the ubiquitin ligase activity associated with CUL2^ZER1^ is required for the growth-promoting activity of HPV16 E7. However, we have not ruled out the possibility that an activity of ZER1 unrelated to CUL2 is essential in HPV-positive cells. Future work will focus on determining whether the ligase activity of CUL2^ZER1^ is required for the growth-promoting activity of HPV16 E7, and if so, what substrates are targeted for ubiquitination by HPV16 E7/CUL2^ZER1^. Given that ZER1 is essential in HPV-transformed cells regardless of HPV genotype, future studies will investigate the role of ZER1 in cancer cells caused by other high-risk HPV. Specifically, we aim to determine whether ZER1 affects the same pathways and/or substrates in HPV16-positive cells as in cells transformed by other high-risk HPVs. Inhibiting ZER1 and/or the HPV16 E7-ZER1 interaction could limit the growth of cancers caused by high-risk HPV.

## MATERIALS AND METHODS

### Plasmids and Cloning

Chimeric HPV16 and 18 E7 genes were cloned using splicing by overlap extension PCR. N-terminal and C-terminal HPV16 E7 and HPV18 E7 sequences were amplified using a pDONR-Kozak-HPV16 E7 plasmid (Supplemental Table 1). HPV16 E7 mutant 1 (R49A, H51A, H73A, E80A, D81A), mutant 2 (I93A, K97A, P98A), HPV16 E7 H73A and E80A/D81A sequences were synthesized by IDT and AttB1 and AttB2 sites were added by PCR. HPV E7 sequences were cloned into the pDONR 223 GAW vector. pDONR HPV E7 sequences were recombined into MSCV-P C-Flag-HA GAW or MSCV-neo C-HA GAW vectors. ZER1 sgRNAs sequences from the Broad Institute Brunello library were cloned into the LentiCRISPR v2 vector following standard protocols (59). A list of all the plasmids used in this paper is in Supplemental Table 2.

### Cell culture and transductions

Primary Human Foreskin Keratinocytes (HFK) were obtained from the Skin Biology and Diseases Resource-based Center (SBDRC) at the University of Pennsylvania. HFK and N/Tert-1 cells (hTert-immortalized HFK) (60) were cultured in keratinocyte serum-free media (K-SFM) supplemented with bovine pituitary extract, EGF, CaCl_2_ and penicillin/streptomycin. SiHa, HeLa, Caski and C33A cells were cultured in Dulbecco’s Modified Eagle Medium (DMEM) supplemented with 10% fetal bovine serum and penicillin/streptomycin. During cycloheximide treatments, cells were treated with 40 µg/ml cycloheximide in cell culture media and harvested at the indicated time points.

Retroviral and lentiviral constructs were packaged in 293 Phoenix or 293T cells respectively as described in (61). HFK, N/Tert-1, SiHa, Caski, HeLa, and C33A cells were transduced with retroviruses to stably express GFP or an empty vector as controls, or with retroviruses encoding HPV16 or HPV18 E7 as previously described (8). HFK, N/Tert-1, SiHa, HeLa, and C33A cells were engineered with LentiCRISPRv2 lentiviruses encoding spCas9 and sgRNAs targeting ZER1, PTPN14, or non-targeting controls as previously described (58). One day after transduction, engineered keratinocytes and cervical cancer cell lines were selected with hygromycin, puromycin, or G418 alone or in combination.

For immunoprecipitation experiments, N/Tert-1 cells were grown at high density using a 1:1 mixture of K-SFM and DF-K media (DF-K mix) as previously described (8). Briefly, cells were grown to 30% confluency, after which the media was replaced every other day with DF-K mix. Cells were harvested at around 85% confluency and pellets were flash-frozen in liquid nitrogen or dry-ice with 100% ethanol. Pellets were stored at −80oC until ready for processing or processed immediately. For immunoprecipitation experiments, one 15-cm plate was used for each engineered N/Tert-1 cell line. Six 15-cm plates of N/Tert-1 16E7 cells were used for each gel filtration chromatography experiment.

### Lifespan extension assay

Primary HFK were engineered and cultured as described in cell culture and transductions. Population doublings were calculated based on the number of cells collected and plated at every passage. The growth of the cells was monitored for one to two months.

### Western blots, antibodies, and immunoprecipitation

Western blots were performed by separating protein lysates on 4-20% gradient polyacrylamide gels (Bio-Rad) and transferring protein to polyvinylidene difluoride (PVDF) membranes as previously described (8). Membranes were blocked in 5% nonfat dried milk in TBST (Tris-buffered saline pH 7.4 with 0.05% Tween 20) and then incubated with primary antibodies to ZER1 (GeneTex), CUL2 (Bethyl), RB1 (Calbiochem/EMD), UBR4 (gift of Yoshihiro Nakatani), GAPDH (Invitrogen), Actin (Millipore), HA-HRP (Roche), or PTPN14 (Cell Signaling Technology). Antibody information is in Supplemental Table 3. Membranes were washed in TBST and incubated with a horseradish peroxidase (HRP)-linked anti-mouse or anti-rabbit secondary antibody and detected using chemiluminescent substrate. For anti-HA immunoprecipitations, HA-tagged proteins were immunoprecipitated with anti-HA agarose beads (Sigma) and processed for Western blotting as previously described.

### Gel filtration chromatography

Fresh N/Tert-1 cell pellets were lysed in mammalian cell lysis buffer (MCLB) [50mM Tris pH 7.8, 150mM NaCl and 0.5% NP40] containing protease and phosphatase inhibitors at a final concentration of: 10mM sodium fluoride, 0.1mM sodium orthovanadate, 5mM b-glycerophosphate, 0.5mM phenylmethylsulfonyl fluoride, 5ug/mL leupeptine and 1ug/mL pepstatin A. The cell lysate was incubated for 15 minutes on ice and then centrifuged at maximum speed for 10 minutes. The supernatant was further filtered using a 0.2um PVDF syringe filter (Cytiva). The Superose 6 10/300-Gl column (Cytiva) was loaded with about 6 mg of total protein and run on a Pure fast protein liquid chromatography (FPLC) with a flow rate of 0.5ml/min. Five-hundred microliter fractions were collected. The even numbered fractions were subjected to HA-immunoprecipitation and analyzed by Western blot. Size standards were generated using a gel filtration marker kit of protein molecular weights that ranges from 29,000 to 700,000 (Sigma-Aldrich).

### RNA sequencing

The RNeasy mini kit (Qiagen) was used to isolate total RNA from three independently transduced populations of HFK-GFP, HFK-16E7, or HFK 16E7 E80A/D81A cells. PolyA selection, reverse transcription, library construction, sequencing, and initial analysis were conducted by Novogene. Relative expression data for selected genes was extracted using Novosmart analysis software.

### DepMap dataset

ZER1 chronos dependency scores were downloaded from the DepMap website (https://depmap.org/). The dataset was graphed based on the type of cancer and the HPV status.

### Anchorage-independent growth assay

SiHa, HeLa, and C33A cells were engineered as described above in cell culture and transductions. One passage after drug selection was completed, 2.5×10^4^ cells were mixed with 0.4% Noble agar and seeded into 6-cm plates that were coated with 0.6% Noble agar, both in DMEM supplemented with 10% fetal bovine serum and antibiotic/antimycotic. Cell lines were seeded in triplicate plates for each condition tested and colonies formed for about 3 weeks. Images of soft agar plates were taken using a Bio-Rad Gel Doc imager. Colonies were counted using ImageJ.

## ACKNOWLEDGEMENTS

We thank the members of our laboratory for helpful discussions. We thank Ronen Marmorstein, Ph.D., and members of his laboratory Shirley Zeng and Karen Zhang for their help running FPLC. This work was supported by American Cancer Society grant 131661-RSG-18-048-01-MPC and NIH/NIAID R01 AI148431 to EAW and by NIH/NIAID R01 AI148431-S1 to JN and EAW.

## FIGURE LEGENDS

**Supplemental figure 1** | **HPV16 E7 E80A/D81A prolongs the half-life of RB1**. Quantification of RB1 protein levels in the cycloheximide experiment. The intensity of the bands in the RB1 western blots was quantified using ImageJ software. The graph indicates the amount of RB1 protein at each time point after cycloheximide treatment, relative to the empty vector control at t=0h. The half-life of RB1 (t_1/2_) was calculated for each condition based on a linear regression of each data set.

**Supplemental figure 2** | **HPV-positive cells depend on ZER1 to survive**. ZER1 chronos dependency scores of multiple cancer cell lines were downloaded from the DepMap website (https://depmap.org/). Each dot represents a cell line. A score of 0 denotes that a gene is nonessential whereas a score of −1 corresponds to the median of all common essential genes. All but two of the cell lines with Chronos scores ≤ −0.5 contain HPV DNA.

